# In Vivo Production of a Protein via the Delivery of Plasmid DNA by Computationally Designed Polymer Nanoparticles (PNP)

**DOI:** 10.1101/2025.02.14.638338

**Authors:** John D. Fisher, Shannon R. Petersen, Olivia Heynes, Suteja Patil, Teresa Tamayo-Mendoza, Felipe Oviedo, Gustavo Guzman, Jared Van Reet, Jeffrey M. Ting, Sean H. Kevlahan, Thomas X. Neenan, Shashi K. Murthy

**Affiliations:** Nanite, Inc., Boston, Massachusetts, USA

## Abstract

While monoclonal antibodies (mAbs) are a prominent class of pharmaceutical products there remain significant issues of cost, complexity, and especially delivery: concepts that overcome the need to infuse antibodies frequently are a desirable objective. An attractive approach is to deliver non-integrating DNA directly to muscle tissue, allowing the patient to act as their own so-called “protein factory.” Demonstrations of this concept have been made using lipid nanoparticles (LNPs) and viral vectors, but these delivery methods face significant challenges, including poor extrahepatic delivery, cargo compatibility, safety, redosability, and cost. Polymer nanoparticles (PNPs) offer a solution to these problems, but face their own challenges, such as the vast number of possible polymer structures, and polyplex formulation conditions. However, advances in machine learning, materials informatics, and high-throughput chemical synthesis techniques provide a foundation for addressing these challenges by efficiently exploring the polymer design space. Our SAYER^TM^ platform utilizes a large computational dataset of plasmid DNA (pDNA) - polymer interactions to facilitate targeting agent discovery and refinement via deep learning, and to drive discovery of novel PNPs for a wide variety of target tissues. In this work, we demonstrate our ability to design PNPs that can deliver pDNA encoding for PGT121, a broadly neutralizing anti-HIV antibody that targets a V3 glycan-dependent epitope site on the HIV-1 envelope glycoprotein. SAYER-designed polymers formed small and stable PNPs with PGT121 plasmids. Intravenous delivery demonstrated strong serum PGT121 protein levels at 1-day post-transfection and higher levels of protein expression when compared to other state-of-the-art DNA-delivery vehicles. More importantly, intramuscular delivery of Nanite PNPs enabled greater than 1.0 µg/mL peak protein expression levels, with meaningful, durable expression levels at > 56 days post-injection. Further, we showed that we can increase antibody levels and durability by redosing. A general trend of lower dose and lower N/P ratio providing higher antibody levels is seen when PNPs were delivered intramuscularly. These parameters, separate from polymer structure, provide different mechanisms to optimize PNP in vivo delivery performance using machine learning techniques. Extension of the concept to the continuous production of other antibodies, proteins or enzymes is possible, suggesting the broad applicability of pDNA depoting via PNPs as a therapeutic modality. Finally, we emphasize that this strategy of delivering a DNA-encoded secreted proteins via a safe and effective PNP in vivo could be applicable to a wide range of other disease modalities.

## Introduction

The importance of biologics (antibodies, bispecifics, peptides, and oligonucleotides) as therapeutics continues to grow, with estimates that the biologics market will exceed that of small molecules by 2027. Within the biologics market, antibodies and related entities such as bispecifics or drug-antibody conjugates play an outsized role, with >100 monoclonal antibodies (mAbs) approved for a range of clinical applications, including immunology, oncology, and hematology. mAbs have now achieved the status of best-selling pharmaceutical products: as of 2024, eight out of the ten bestselling drugs globally were mAbs; and in 2023 one single mAb, the PD-1 inhibitor, Keytruda (pembrolizumab) generated sales of $25 billion. However, routes of administration and frequency of dosing remains an issue with these therapies, with many of the most popular antibodies on the market requiring dosing on weekly or bimonthly schedules to achieve serum levels in the µg/mL range. While most antibodies are still delivered via intravenous (i.v.) infusion in a healthcare setting, subcutaneous (s.c.) injection as a delivery modality continues to grow. Compared with the i.v. route, the s.c. delivery route is less invasive, faster to administer, and, increasingly allows for self-administration in a home setting. Conventionally, it is accepted that the maximum volume tolerable via a single s.c. injection is ∼1.5-2.0 mL. Aggregation of antibodies at high concentrations can increase viscosity and decrease solution stability, thus limiting clinically acceptable concentrations to below 150-200 mg/mL.^1^ With these constraints, there remain challenges in the effective delivery of the large loading doses required for many therapeutic applications. These factors, together with the frequent requirement for patients to travel to specialized centers for i.v. infusions, with attendant time and productivity costs, may hamper the full deployment of otherwise life-saving medications. In recent years, the broad field of mAbs has seen many advances to overcome these challenges, while simultaneously broadening their reach across disease areas. Computationally driven design of mAbs represent an example of new approaches. However new modalities that overcome the delivery challenges of antibodies remain a desirable objective.

An attractive approach is to deliver DNA or mRNA encoding for the antibody of interest directly to patients. This concept allows the patient to act as their own so-called “protein factory,” producing the therapeutic mAb for prolonged periods of time, and to secrete it, either systemically or locally, depending on the route of administration and the tissue of residence. The protein factory concept in principle offers a convenient alternative to the conventional production, purification and administration of mAb proteins. Prolonged in vivo production of mAbs may contribute to (i) a broader adoption of mAbs in price-sensitive markets, (ii) improved accessibility to therapy in both developed and developing countries, and (iii) more effective and affordable treatment modalities, e.g. by facilitating nucleotide-based mAb cocktails or local mAb expression.^2^

While the use of mRNA has risen in recent years,^3^ plasmid DNA (pDNA) is generally cheaper to produce, ship, and store, and has a much longer shelf life than mRNA. After entry into the nucleus, pDNA remains in a non-replicating, non-integrating episomal state, and is only lost during the breakdown of the nuclear envelope at mitosis. Additionally, pDNA has no defined restrictions regarding the size of the transgene, and its modular nature allows for straightforward molecular cloning, making it easy to manipulate and design for therapeutic use. There is ample precedent for the use of pDNA in ongoing or completed gene therapy clinical trials where it has been shown to be well-tolerated and safe. Successful delivery of pDNA to elicit mAb production would therefore appear to be a viable strategy to develop innovative biologics therapeutic platforms.^4^

### Delivery of pDNA

Delivery of pDNA to cells for protein vectorization has many of the same delivery challenges presented by conventional gene therapies such as gene or base editing.^5^ Traditionally delivery of genetic cargoes has been accomplished via the use of viral vectors or lipid nanoparticles (LNPs). Evolutionally, viral vectors have evolved to release their genome into target cells and are therefore ripe for further engineering for a variety of purposes, including genetic cargo delivery and vaccines. Surface proteins of viruses are typically capable of overcoming cellular physical barriers and nuclear localization, thus allowing the efficient deposition of their cargo (i.e., mRNA, DNA, mixtures and other materials) into the target cell. A variety of types of recombinant viral vectors have been used in the gene therapy field including lentivirus (LV), gamma retrovirus (γ-RV), adenovirus (AdV), and adeno-associated virus (AAV). Viral vectors are effective in the delivery of DNA but despite the great clinical success of AAV-based gene therapies, the cargo capacity of these vectors is generally limited to applications using expression cassettes no larger than ∼4.75 kb. In addition, despite substantial advancements in the field of viral capsid engineering, re-dosing of viral vectors remains problematic and, in the context of in vivo generation of mAbs that may require frequent dosing over the life span of the patient, highly impractical.

In principle, LNPs are an attractive option to viral delivery for pDNA. The 2018 approval of Onpattro (Patisiran) for hereditary transthyretin amyloidosis established the concept of LNPs as a viable approach to genetic cargo delivery. LNPs played a pivotal role in the global COVID-19 vaccination campaigns, marking a significant milestone and highlighting their potential as potential gene delivery agents. Despite the success of LNP-based therapeutics in the COVID era, this approach faces several challenges including poor tissue tropism outside of the liver, efficacy of delivery, and potential side-effects including toxicity and immunogenicity. LNPs are composed of mixtures of (generally 4 or 5) ionizable lipids, helper lipids, polyethylene glycol (PEG)-lipids and cholesterol, each serving specific functions. LNPs primarily accumulate within the liver after i.v. administration. Efforts to enhance the efficacy and biodistribution of LNPs by modifying their lipid composition have received extensive attention in recent years. However, the physical and chemical constraints posed by the thermodynamic challenge of forming a lipid nanoparticle greatly restrict the diversity space from which new LNPs can be designed. Additionally, the worldwide deployment of LNPs in COVID vaccine, have led to high levels of antibody responses to polyethylene glycol (PEG)-lipids, thus hampering the use of this critical LNP component. Further, the production of LNP formulated genetic cargoes requires specialized production equipment and the resulting particles tend to have poor stability and limited shelf life. Significantly, in the current context it is well established that pDNA-LNPs have poor transfection efficiency, are highly inflammatory and induce mortality at commonly used therapeutic doses in naïve mice. Various mechanisms for this toxicity have been proposed including the inappropriate activation of the cGAS-STING signaling pathway.^6^

With the limitations of both viral vectors and LNPs as delivery agents, there is therefore a recognized need to develop more robust nanocarriers that can achieve DNA delivery with high specificity, reliability, and safety. Polymer nanoparticles (PNPs), which are polyplexes formed from tailored cationic polymers and therapeutic nucleic acids, offer distinct advantages as multifunctional delivery systems and may be ideally suited for the protein factory concept. Advanced polymer chemistry techniques allow access to a vast variety of conceivable chain structures and architectures from a diverse array of building blocks, and the resulting polymeric materials can be readily synthesized at scale, can be designed to be excreted or absorbed, and form thermodynamically stable particles. Additionally, there is an excellent track record in the use of polymer materials as drug delivery agents, excipients and as structural materials in advanced medical devices.^7^ While the utility of polymers for drug delivery is a well-established concept for small molecules, therapeutic biologics present distinct challenges. DNA, RNA, and genomic editing ribonucleoproteins are larger, hydrophilic, ionic, and prone to degradation. Prospective polymer delivery systems for DNA delivery need to balance opposing attributes for these payloads by providing (i) colloidal stabilization across multiple biological barriers, and (ii) efficient payload release at the site of action. While cationic polymers have less clinical experience for the delivery of genetic drugs compared to viral vectors or lipid nanoparticles, they have emerged as promising candidates for the delivery of biologic agents due to their wide possible chemical space, which allows for the discovery of materials with properties such as cell and tissue tropism and capacity for endosomal escape. In the presence of water, these polymers undergo hydrolytic degradation into byproducts that are either resorbable, or benign and small enough for renal clearance.

Despite these inherent advantages, there remains a major limitation to the wider adoption of polymers as gene therapy agents, due in part to the lack of systems and methodologies to explore the vast landscape of the potential polymer design space. Criteria such as polymer architecture, molecular weight, monomer diversity etc. lead to estimates of the size of the potential polymer landscape to be on the order of 10^60^— a vast number that, even if reduced to more surmountable subsets via exhaustive data mining, cannot be practically interrogated with the current state of polymer informatics.^8^ Secondly, in vitro cellular assays, while amenable to high-throughput screening for highly valuable transfection data across cell types, often provide poor predictions on the crossing of complex physiological barriers in vivo Thus, unclear in vitro–in vivo correlation results represent a difficult obstacle in PNP optimization in translating genetic drugs from cellular level assays to living systems.^8^

Nanite’s proprietary platform, SAYER, offers a comprehensive solution to these challenges by integrating experimental data generation with artificial intelligence (AI) techniques including machine learning and cheminformatics. This integration enables the navigation of the polymer design space with unprecedented efficiency and precision, identifying high-performing PNPs for the efficient delivery of diverse genetic cargo to tissues outside of the liver.^9^ By leveraging curated, high-quality experimental and synthetic data, SAYER^TM^ transforms the historically fragmented and unstructured polymer research landscape into a machine-actionable framework. Its capability to handle high-dimensional datasets allows researchers to model complex structure-function relationships, facilitating the discovery of polymers optimized for specific cargos.^8^ Additionally, SAYER’s adaptive learning models iteratively improve with every data point, ensuring increasingly accurate predictions of polymer performance across both in vitro and in vivo conditions (Fig. 1). This increase in prediction accuracy reduces the number of candidates needing to be tested experimentally while at the same time allowing systematic exploration of new chemical design spaces.

**Figure 1.**
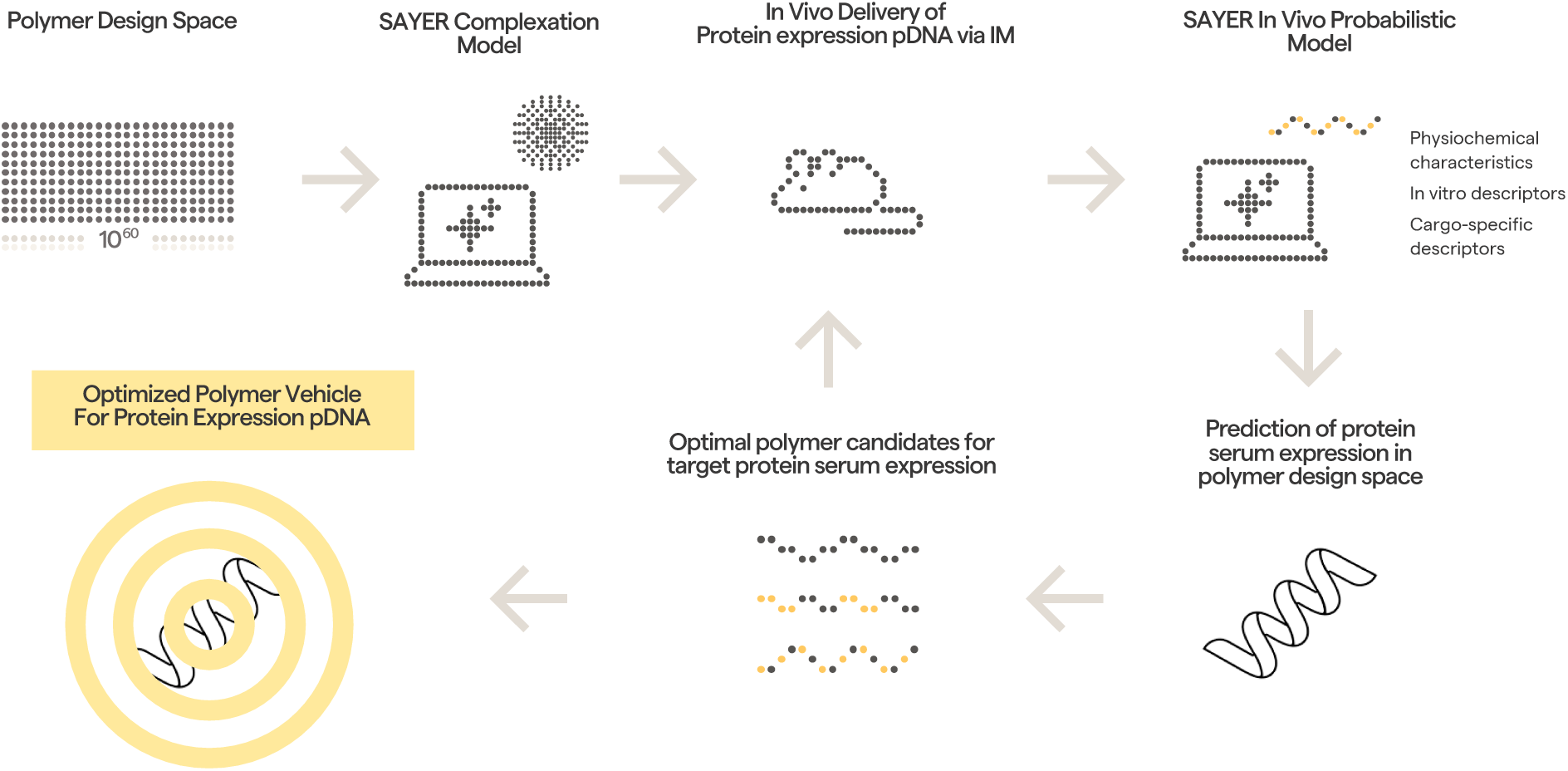
Illustration of the SAYER platform as applied to the use case of protein expression via delivery of pDNA payloads using polymer nanoparticles.

We have previously described our success in developing polymeric nanoparticles that are highly tropic to the lung based upon the insights from our SAYER platform.^8^ Additionally, we have recently reported our results on a PNP-based in vivo CAR-T CD19 construct that shows robust tumor killing in humanized mice as well as expression in primates with minimal toxicity.^10^ These projects, together with other undisclosed programs suggest the broad applicability of polymer nanoparticles as delivery agents for genetic cargoes.

### Vectorization of antibodies using a polymer nanoparticle

In the present work, we demonstrate our ability to design PNPs that can deliver pDNA encoding for PGT121,^11^ a broadly neutralizing anti-HIV antibody, at high expression levels and with high antibody persistence doses as long as 56 days. PGT121 is a recombinant human IgG1 monoclonal antibody that targets a V3 glycan-dependent epitope site on the HIV-1 envelope glycoprotein. Structural studies suggest that PGT121 inhibits the binding of gp120 to the CD4 receptor by interfering with Env receptor engagement and blocking viral entry. It has been reported that PGT121 decreases viral load in simian–human immunodeficiency virus (SHIV)-infected rhesus macaques. Additionally, PGT121 delays SHIV rebound when combined with a Toll-like receptor 7 agonist and protected uninfected monkeys against SHIV challenge at a plasma concentration of <5 μg/mL. A series of Phase 1 and Phase 2 studies^12^ have been completed with PGT121 focused on safety, tolerability, pharmacokinetics, and antiviral activity. Studies to date suggest that PGT121 is generally safe and well tolerated. The in vivo production of PGT121 by viral vectors has been demonstrated.^13^ However, a major challenge with this approach is the inability to redose and the high cost of production.^14^

The kinetics of PGT121 have further been examined in at least three trials.^15^ A two-part Phase 1 trial (NCT02960581) initially evaluated a single i.v. or s.c. dose (3, 10, or 30 mg/kg) of PGT121 in adults without HIV and with HIV on anti-retroviral therapy (ART), followed by a single i.v. dose (30 mg/kg) of PGT121 in participants with HIV. The elimination half-life of PGT121 was 22 days in participants without HIV, 16 days in participants with HIV and on ART, and 14 days in participants with HIV who were both viremic and off ART. In a Phase 1/2a trial (IAVI T003; NCT03721510) evaluating triple bNAb therapy with PGT121 and two other antibodies, PGDM1400 and VRC07-523LS, administered via i.v. infusion in adults with and without HIV, the median elimination half-life of PGT121 was approximately 19.9 days. In a Phase 1 study (NCT04212091) of i.v. or s.c. infusions of PGT121.414.LS (a PGT121 variant with a mutation designed to extend in vivo half life) in healthy adults without HIV, PGT121.414.LS demonstrated a half-life ranging from 53.6 to 74.3 days.

In the current work, we demonstrate that the kinetics of PGT121 can be dramatically extended using our PNP/pDNA approach. In addition, we demonstrate that re-dosing is viable and that expression levels of PGT121 can be maintained at µg/mL levels for up to 56 days in mice when the pDNA is delivered to skeletal muscle. No evidence of local or systemic toxicity was observed, suggesting that this approach may offer a viable route for the development of long-lasting HIV treatments. Extension of the concept to the continuous production of other antibodies, proteins or enzymes is possible, suggesting the broad applicability of pDNA depoting as a therapeutic modality.

## Experimental Section

### Materials & Methods

#### Polymer Synthesis

A family of cationic polyesters designed for the delivery of PGT121 was prepared using insights from our computational and high throughput synthetic capabilities. The polymers were prepared using a combination of ring-opening copolymerization (ROCOP) and thiol-ene “click” reactions. ROCOP of epoxides and anhydrides produces alternating copolymers and is a widely used, controlled polymerization technique for the synthesis of polyesters. Scaffold polymers functional for future modifications using thiol-ene chemistry were prepared by using monomers containing alkene functionalities. The alkenes were present on either the epoxide, anhydride, or both. Once prepared, the candidate scaffold polymers were divided and functionalized with a combination of cationic thiols and either neutral or anionic thiols to produce a large number of samples for study. A protypical structure of shown in Fig. 2. In the current work, as detailed below, four polymers from this class (D5588, D3503, D6352, and D5586) were identified using our machine learning approach. Structures are not disclosed in this publication.

**Figure 2.**
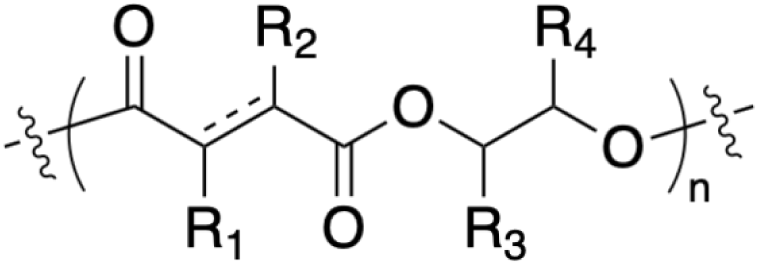
Representative structure of a Nanite polymer utilized for pDNA delivery.

#### Machine Learning Approach

Machine learning-driven polymer optimization aims to drastically reduce the number of failed synthesis and testing cycles required before identifying a polymer nanoparticle with the desired cargo binding capacity, transfection efficiency, biocompatibility, and stability. By leveraging SAYER’s predictive ML algorithms, we dramatically increase the likelihood that a newly synthesized polymer exhibits the necessary characteristics for effective gene delivery. This effectively raises the probability of success for each experiment, thereby reducing the total number of experimental iterations required to achieve a viable polymeric delivery vehicle.

In vivo experiments for DNA delivery are inherently slow and resource-intensive, necessitating a strategy that maximizes the probability of success for any polymer tested in an animal model. Given the vast chemical space of potential polymers, we aimed to dramatically down sample candidates at each stage of the experimental campaign, ensuring that the few polymers advancing to in vivo studies had the highest likelihood of achieving efficient transfection. First, we identified a relevant subset of polymer families from the extensive chemical databases available in SAYER, reducing the candidate pool from an astronomical ∼10⁶⁰ possibilities to several thousand. Next, a machine learning model trained using our proprietary data set of binding interactions between polymer and payload further refined this set to a few hundred candidates. Subsequently, a second ML model, trained on data from in-vitro transfection experiments, selected a few dozen polymers with high transfection efficiency in target cells. This multi-stage selection process is shown in Fig. 3

**Figure 3.**
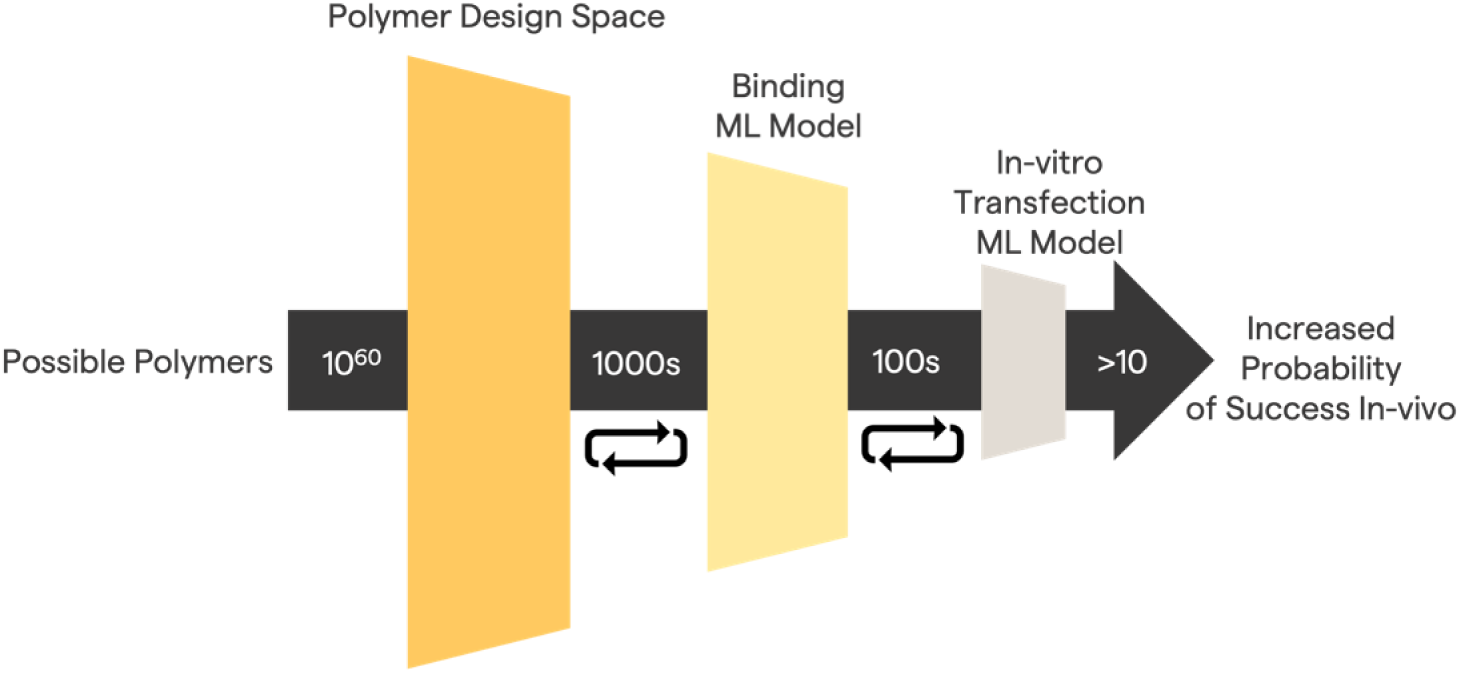
Multi-stage section process. Our ML models included in the Sayer platform allow us to quickly explore and down select polymers reducing the experimental iterations, especially in low-throughput and costly experiments.

#### DNA Cargos

ZsGreen plasmid (3.6 kb) was purchased from VectorBuilder (Chicago, IL) by cloning the ZsGreen1 coding sequence into a CMV enhancer/promoter and SV40 polyA expression vector. PGT121 nanoplasmid (3.6 kb) was purchased from GenScript (Piscataway, NJ) by cloning the full PGT121 coding sequence into a GenCircle dsDNA vector. The CMV enhancer/promoter drives expression of both the heavy chain (no stop codon) and the light chain of PGT121, that are separated by a furin cleavage site upstream of a P2A linker. Downstream of the light chain stop codon is a bGH polyA tail. After the PGT121 fusion protein is expressed in a cell, cleavage of the furin and P2A sites occurs, producing a single PGT121 heavy chain and a single PGT121 light chain.

#### Dynamic Light Scattering (DLS)

PNPs used for DLS analysis were prepared using an automated OT-2 liquid handler robot from Opentrons Labworks Inc. (Long Island City, NY). Briefly, PNPs were formed by adding 1 volume of polymer at the specified N/P ratio to 1 volume of PGT121 nanoplasmid followed by gentle mixing via pipetting. Here, N/P ratio represent the ratio of the positively charged nitrogren (N) groups from the cationic polymer to the negatively charged phosphate (P) groups from the backbone of the plasmid DNA. All steps were performed at room temperature in 5% glucose solution. Approximately 30-45 mins after formulation, PNP samples were read on an automated DynaPro Plate Reader III from Wyatt Instruments (Santa Barbara, CA) with an acquisition time of 1 second and 5 acquisitions per well. Autocorrelation functions on DYNAMICS software from Wyatt Instruments (Santa Barbara, CA) were used to calculate the mean hydrodynamic radius and polydispersity of each condition. Reported values were calculated using regularization (plots) and cumulant fit (table) algorithms. n=1 read per condition.

#### In Vitro Transfections

HEK293T cells were cultured in DMEM media supplemented with 10% fetal bovine serum (FBS) and 1X penicilin-streptomycin in a Forma Direct Heat CO2 incubator from ThermoScientific (Waltham, MA), set at 37 °C and 5% CO2. Approximately 24 hrs before transfection, ∼6,500 HEK293T cells were seeded (DMEM + 10% FBS) in each well of a poly-L-Lysine coated 96 well plate. Using an automated OT-2 liquid handler robot from Opentrons Labworks Inc. (Long Island City, NY), PNPs were formulated by adding 1 volume of polymer at the specified N/P ratio to 1 volume of DNA cargo followed by gentle mixing via pipetting. All steps were performed at room temperature in 0.9% NaCl solution. Approximately 45 minutes after PNP complexation, 50 µL of PNP solution containing 250 ng of DNA cargo was added to each well of the 96 well plate and returned to the incubator. For ZsGreen plasmid experiments, 72 hours after transfection, cells were stained with Hoechst 33342 and media was replaced with FluoroBrite DMEM supplemented with GlutaMAX and live-cell imaged on an Operetta CLS high-throughput confocal imager from Revvity (Waltham, MA). The effective transfection efficiency was calculated by taking the product of the transfection efficiency (% ZsGreen+ cells) and the viability (% total cells relative to negative control) per well. Heat maps report the effective transfection efficiency and SEM from n=2 experiments. For PGT121 nanoplasmid experiments, 6-7 days after transfection, supernatant was collected from each well, spun down at 1,000 g for 5 mins to remove cell debris, flash frozen in liquid nitrogen, and placed at –80 °C until samples were run on ELISA, with n=2 experiments per condition.

#### Animal Work

Animal experiments were performed by Biomere Biomedical Research Models, Inc. (Worcester, MA). All animal work was carried out in accordance with NIH Office of Laboratory Animal Welfare (OLAW) and Association for Assessment and Accreditation of Laboratory Animal Care (AAALAC) assurance. For all animal experiments, approximately 7-week-old female RAG2-/-mice from Taconic Biosciences, Inc. (Germantown, NY) were administered PNPs at Day 0. PNPs were formed by adding 1 volume of polymer at the specified N/P ratio to 1 volume of PGT121 nanoplasmid followed by gently mixing via pipetting. All steps were performed at room temperature in 5% glucose solution. Approximately 45 mins after formulation, PNPs were injected into mice. For intravenous administration experiments, 1.8 mg/kg of PGT121 nanoplasmid-containing PNP was injected via the lateral tail vein, following a standard protocol and monitored over time. Serial blood draws were taken on days 1, 4, 7, 14, 21, and 28, processed to serum, flash frozen, and stored at –80C until samples were run on ELISA. For intramuscular (i.m.) administration experiments, PGT121 nanoplasmid-containing PNP was injected into the right tibialis anterior and left gastrocnemius muscles. The two redosing groups received a second PNP dose on day 8, identical to the first dose on day 0 (4.0 mg/kg PGT121 nanoplasmid total). Serial blood draws were taken on days 1, 4, 8, 14, 21, 28, 35, 42, 49, and 56, processed to serum, flash frozen, and stored at –80 °C until samples were run on ELISA.

#### Enzyme-linked Immunosorbent Assay (ELISA)

For all ELISA experiments, frozen cell culture supernatant or mouse serum samples were thawed on ice for 30 mins, gently mixed, thawed at room temperature for 30 mins, spun down at 2,000 g for 5 mins, serially diluted in ELISA assay buffer, and then added to the ELISA plate. For HEK293T supernatant samples, the Human IgG ELISA Kit (Colorometric) from Bio-Techne (Minneapolis, MN) was used following the manufacturer’s protocols. For mouse serum sample experiments, the SimpleStep anti-hIgG ELISA from Abcam (Waltham, MA) was used following the manufacturer’s protocols. Immediately following the addition of stop solution, ELISA plates were placed in a SpectraMax ID3 plate reader from Molecular Devices (San Jose, CA) and the absorbance at 450 nm for each well was recorded. Raw absorbance values for hIgG standards and samples were then imported into GraphPad Prism software (Irvine, CA). A standard curve for each ELISA plate was generated using a four parameter logistic curve and sample well absorbance values were interpolated, with n=2 for all cell culture supernatant experiments and error bars representing SEM, and n=2 for all mouse serum sample experiments and error bars representing SEM.

## Results

### SAYER optimization of PNPs for DNA delivery

In this study, we develop first-principles data pipelines and machine learning models for polymer design, leveraging curated molecular representations tailored for predictive accuracy. To incorporate the information on both molecular composition and sequence distribution, we employed a molar-weighted summation of the monomer’s feature vectors, as proposed by Tao et al.^16^ (Fig. 4A). These feature vectors were in turn calculated using modified Morgan Fingerprints.

**Figure 4.**
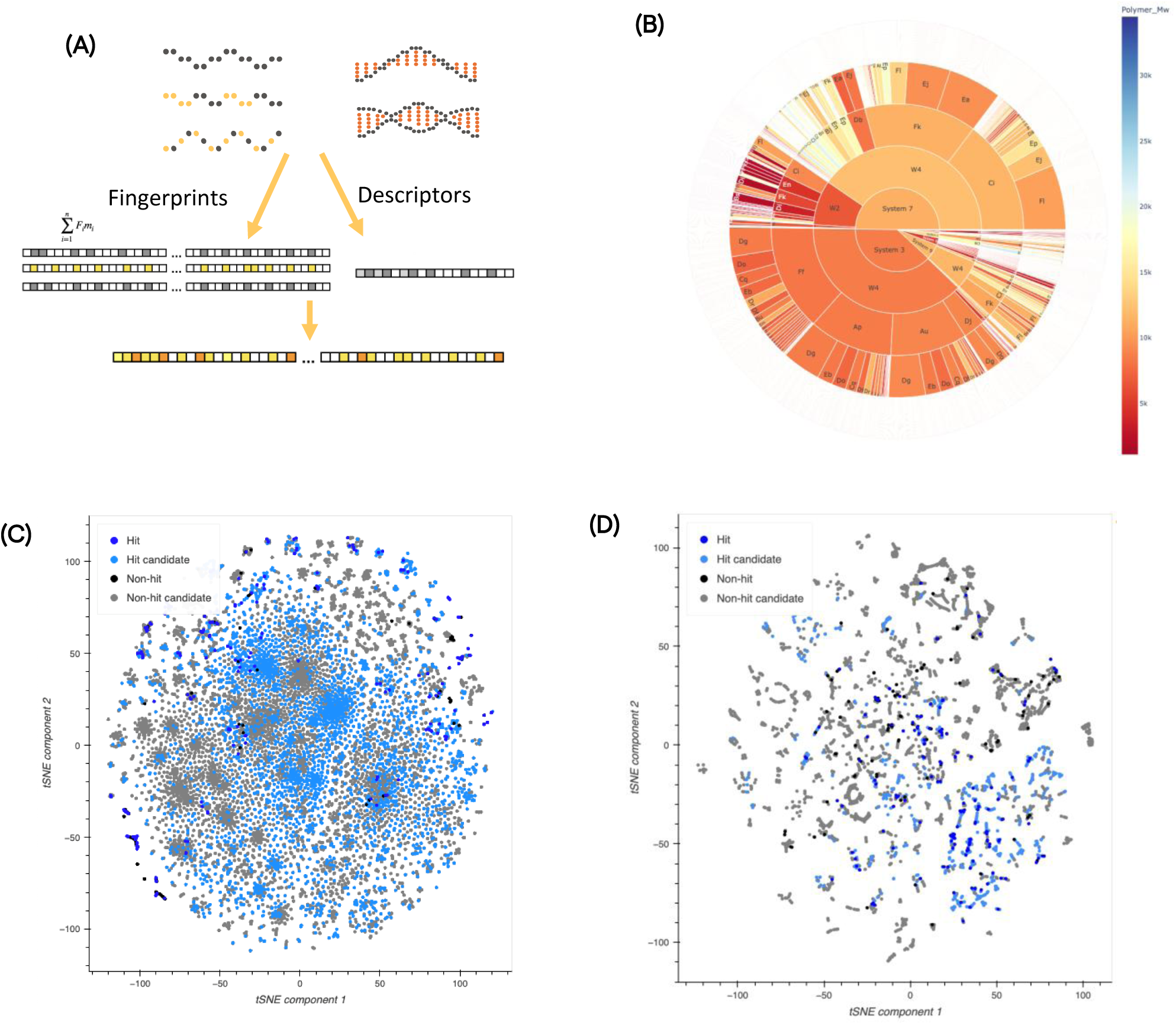
SAYER platform down sample strategy. (A) Polymer representations using molar weighted fingerprints and molecular descriptors. (B) Polymer design space in SAYER platform, where we explored different polymer systems. In the diagram, from inside out each ring displays a different polymer system, different block structure and different monomers. (C) and (D) t-SNE plots using molar weighted fingerprints. (C) Includes polymers screened with binding and solubility experimentally and virtually (candidates) and (D) includes polymers screened for HEK293T pDNA transfection. The hits and non-hits experimentally tested are displayed in dark blue and black respectively, and potential hit and non-hit candidates to explore in future iterations are displayed in light blue and gray. Components 1 and 2 in (C) and (D) represent combinations of multiple variables.

The SAYER platform curates high quality information on a large and diverse set of polymer families (Fig. 4B). To ensure a diverse sampling of the polymer design space, we employ similarity metrics as a quantitative measure of diversity and use it alongside predicted performance to select hit candidates. A key objective is to minimize inefficiencies in experimental synthesis by avoiding insoluble polymers, which we address through a predictive insolubility model. Additionally, a binding model assesses the polymer’s capacity to interact with the target payload (Fig. 4C). By integrating these models, we identify candidate polymers that are both functionally effective and structurally diverse, optimizing the discovery process. After passing these filters, a polymer was further classified as a hit if the mean transfection efficiency in HEK293T cells was predicted to be at least 10% within the range of N/P values tested. Through two iterative selection cycles, we prioritized polymers with high prediction confidence while incorporating a subset of polymer systems underrepresented in existing datasets. This approach expanded both the number and diversity of candidates for in vivo testing while simultaneously improving the predictive performance of our models in in vitro experiments. The polymer space explored in this project is shown in Fig. 4D.

### In vitro validation of PNPs using DNA cargos

We first sought to validate the DNA-delivery efficiency and cytotoxicity of SAYER-optimized PNPs using a ZsGreen reporter plasmid. Polymer stocks were added to a fixed mass of ZsGreen plasmid at varying N/P ratios, the ratio of the positively charged nitrogren (N) groups from the cationic polymer to the negatively charged phosphate (P) groups from the backbone of the plasmid DNA. Simple electrostatic interactions between these molecules allow the PNPs to immediately self-assemble at room temperature. Approximately 45 minutes after PNP formulation, the particles were added to HEK293T cells, a common mammalian cell line used to investigate recombinant protein expression. For ZsGreen protein expression to occur, the PNPs must first enter the cell through electrostatic interactions with the negatively charged cell membrane. Following cellular entry via endocytosis, the PNPs must escape the endosome to avoid degradation within lysosomes. Once the PNPs are released into the cytosol, the plasmid must transport to the nuclear pore complex where it can finally enter the nucleus and begin transcription of the endcoded protein. We observed that all four lead polymers (Fig. 1) efficiently delivered the ZsGreen reporter plasmid in vitro across a range of different N/P ratios (Fig. 5A). For PNPs, there can be a tradeoff between transfection efficiency and toxicity when increasing the N/P ratio of the PNP, and thus the charge of the PNP. We quantified this tradeoff by multiplying the transfection efficiency and the cell viability of the PNP to obtain the effective transfection efficiency (Fig. 5B). All PNPs showed peak effective transfection efficiencies between N/P=3 and N/P=12. These differences in PNP function can be attributed to the structural differences of the polymer chemistries, and thus the structural differences in the PNPs. The observed effects are achieved by varying the properties of the polymer including but not limited to: the composition of cationic monomers, the composition of non-cationic monomers, the relative abundance of each monomer, the physical structure of the polymer (linear vs. branched), and the molecular weight of the polymer. All of the above factors contribute to the physical properties of the PNPs by changing the size, polydispersity, zeta potential (surface charge), and functional delivery capabilities. With even a small but repeatable dataset of in vitro transfection values, polymers (and PNPs) can be tuned to increase delivery efficiency using machine learning techniques. Although these PNPs follow a general trend of increasing transfection efficiency with N/P ratio and eventual toxicity at higher N/P ratios, this is not always the case.

**Figure 5.**
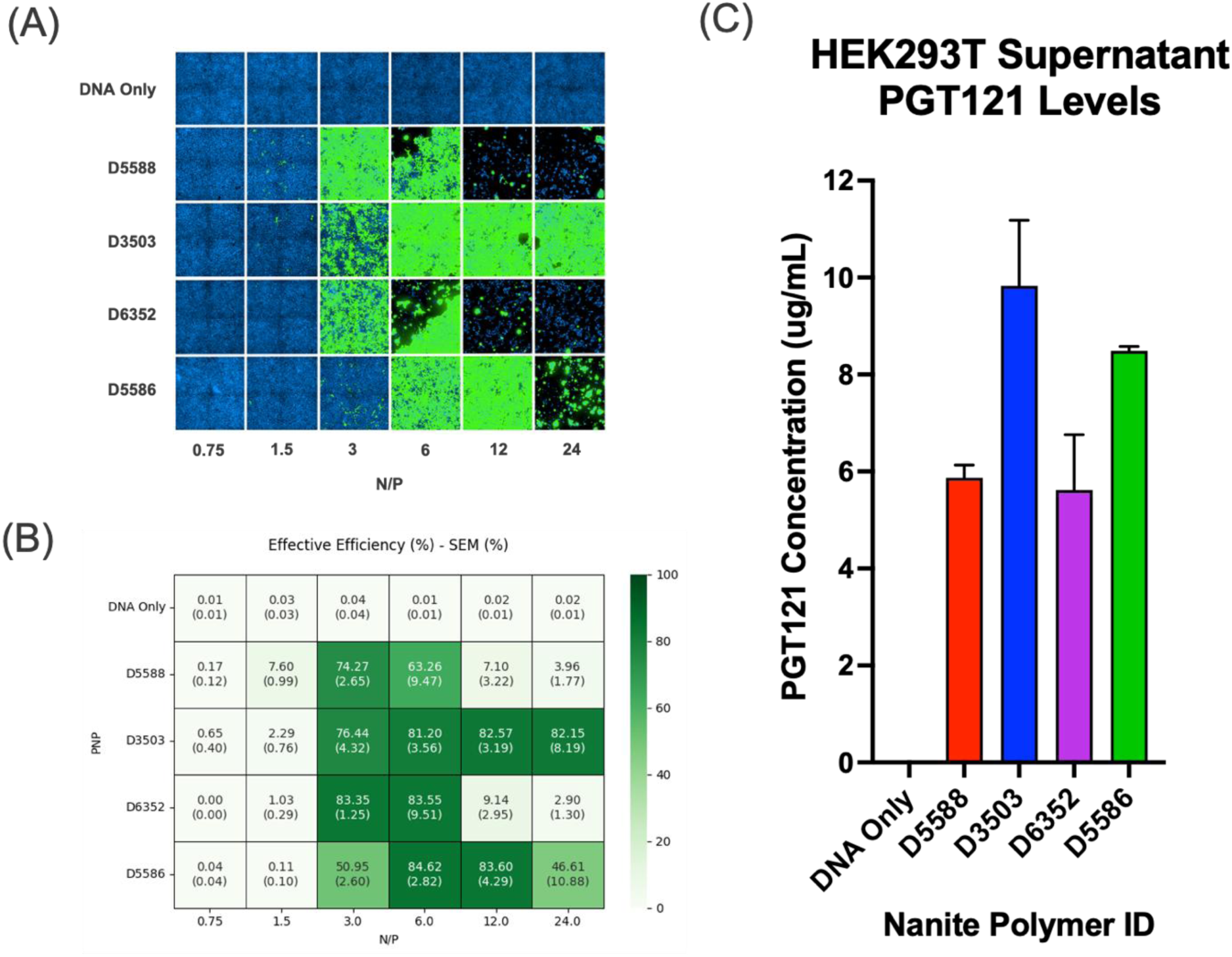
In vitro transfection of Nanite PNPs. (A) Representative images of HEK293T cells transfected with PNPs carrying a ZsGreen reporter plasmid across different N/P ratios. ZsGreen expression and cell nuclei marker are shown in green and blue, respectively. (B) Heat map of the average effective transfection efficiency (product of transfection efficiency and viability) from experiments in (A). (C) HEK293T supernatant PGT121 levels 1 week after in vitro transfection of PGT121 nanoplasmid delivered with Nanite PNPs. n=2-5, error bars=SEM for data in (B), (C).

Following validation that these PNPs could efficiently deliver a cytosolic reporter plasmid, we aimed to determine whether these same polymers could deliver a nanoplasmid encoding the anti-HIV antibody PGT121, a secreted protein. PGT121 is a human IgG (hIgG) antibody which consists of four polypeptides, two identical heavy chains and two identical light chains which self-assemble into a functional antibody molecule via disulfide bonds. To address the need to encode two distinct polypeptides to deliver a functional antibody, we employed a previously developed single plasmid strategy.^17^ Briefly, a nanoplasmid was engineered to encode both the heavy chain and the light chain of PGT121, separated by a self-cleaving P2A linker. Once expressed, the PGT121 fusion protein self-cleaves, forming one heavy chain and one light chain. To validate this PGT121 nanoplasmid, we formulated PNPs and transfected HEK293T cells. Following a period of 6-7 days post-transfection, supernatant was collected and run on an anti-hIgG ELISA to quantify the amount of secreted PGT121 antibody (Fig. 5C). All PNPs produced high levels of secreted PGT121 protein with similar trends to the delivery of the ZsGreen reporter. Due to the similar size of the ZsGreen plasmid (3.6 kb) and PGT121 nanoplasmid (3.6 kb) cargos, this result is not completely unexpected.

After confirming the high delivery efficiency and low toxicity of PNPs carrying the PGT121 nanoplasmid, we sought to confirm the size and uniformity of the particles using dynamic light scattering (DLS). For DLS measurements, PNPs were formulated with PGT121 nanoplasmid in the same manner as the in vitro transfection experiments. A DynaPro plate reader (Wyatt Technology) was used to quantify the size (hydrodynamic radius) and uniformity (polydispersity) of PGT121 PNPs at various N/P ratios (Fig. 6). We observed that these polymers formed small (< 100 nm) and uniform (polydispersity index (PDI) < 0.30) particles that were stable across time (data not shown). Confirmation that we had created small, uniform, and stable PNPs that efficiently deliver PGT121 nanoplasmid without toxicity enabled us to test whether these PNPs could deliver PGT121 in vivo. This stage was also the culmination of the machine learning-guided process which identified the selected polymers shown in Figs. 5-6.

**Figure 6.**
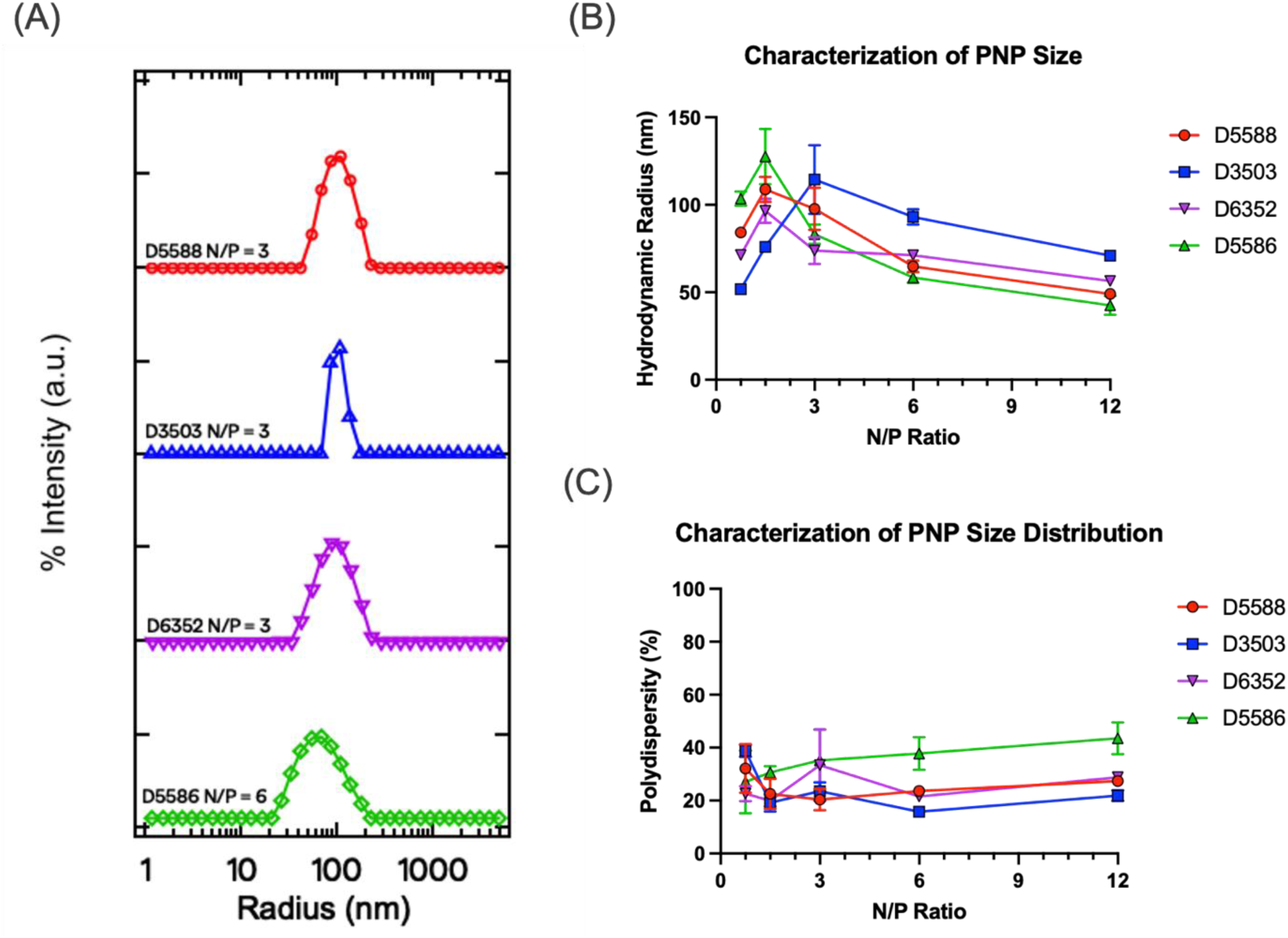
PNP Characterization of PGT121 DNA Cargo by Dynamic Light Scattering. (A) Representative intensity weighted distributions of Nanite PNPs approximately 45 minutes after PNP formulation. (B) Hydrodynamic radius of PNPs across a range of N/P ratios from experiments in Fig. 6A. (C) Polydispersity of PNPs across a range of N/P ratios from experiments in Fig. 6A. n=3, error bars=SEM for data in (B), (C).

### In vivo PNP delivery of a DNA-encoded anti-HIV antibody

Unlike an in vitro environment, there are many physiological barriers PNPs must overcome in vivo to successfully deliver genetic cargo to the nucleus. Different routes of PNP administration in vivo can significantly affect delivery efficiency, just as polymer structure can. To address this issue, we sought to investigate two separate routes of administration commonly used to evaluate gene delivery vehicles in mice: systemic i.v. administration of PNPs into the bloodstream via lateral tail vein injection, and local administration of PNPs to skeletal muscle cells via i.m. injection. When PNPs are injected intravenously, they immediately encounter proteins in the bloodstream which can coat PNPs (the protein corona effect) and influence which organs and cell types they target.^18^ When PNPs are injected intramuscularly, they reach their muscle cell targets immediately, but the intracellular barriers of post-mitotic multi-nuclear skeletal muscle cells pose new obstacles for gene expression.

We first investigated whether the PNPs could deliver PGT121 nanoplasmid intravenously. When administered to a wild-type mouse, human antibodies like PGT121 elicit a predictable immune response. The mouse will produce anti-hIgG antibodies against the foreign human protein, complicating quantitative analysis of gene delivery efficiency. To circumvent this challenge, we utilized the RAG2 knock-out (RAG2-/-) immunodeficient mouse model to carry out in vivo gene delivery experiments. The RAG2 gene is necessary for V(D)J recombination during B and T cell development. RAG2-/-mice produce no mature B or T cells and therefore cannot mount an immune response against exogenous human antibodies. RAG2-/-mice were intravenously injected with PNPs containing 1.8 mg/kg of PGT121 nanoplasmid. Serial blood draws were taken from each animal out to day 28, processed to serum, and run on an anti-hIgG ELISA (Fig. 7A, B). For all PNPs, we observed peak circulating PGT121 expression levels in the 20-100 ng/mL range at 1-day post-injection. We extrapolated an approximate 1-2 week half-life of PGT121 in RAG2-/-mice when nanoplasmid was delivered with our PNPs. These results suggest the PNPs quickly transfected cells in vivo, and that the nanoplasmid was subsequently silenced (CpG silencing) or cleared from the transfected cells. No toxicity was observed (data not shown). We anticipate that optimization of the nanoplasmid promoter/enhancers, 3’ UTRs, codon-optimization, signal peptide, or other genetic engineering could further boost serum PGT121 levels via i.v. PNP administration. Additionally, polymer/PNP optimization to specifically target high-transfection cell types for PGT121 DNA delivery would likely achieve clinically relevant levels of protein antibody expression.

**Figure 7.**
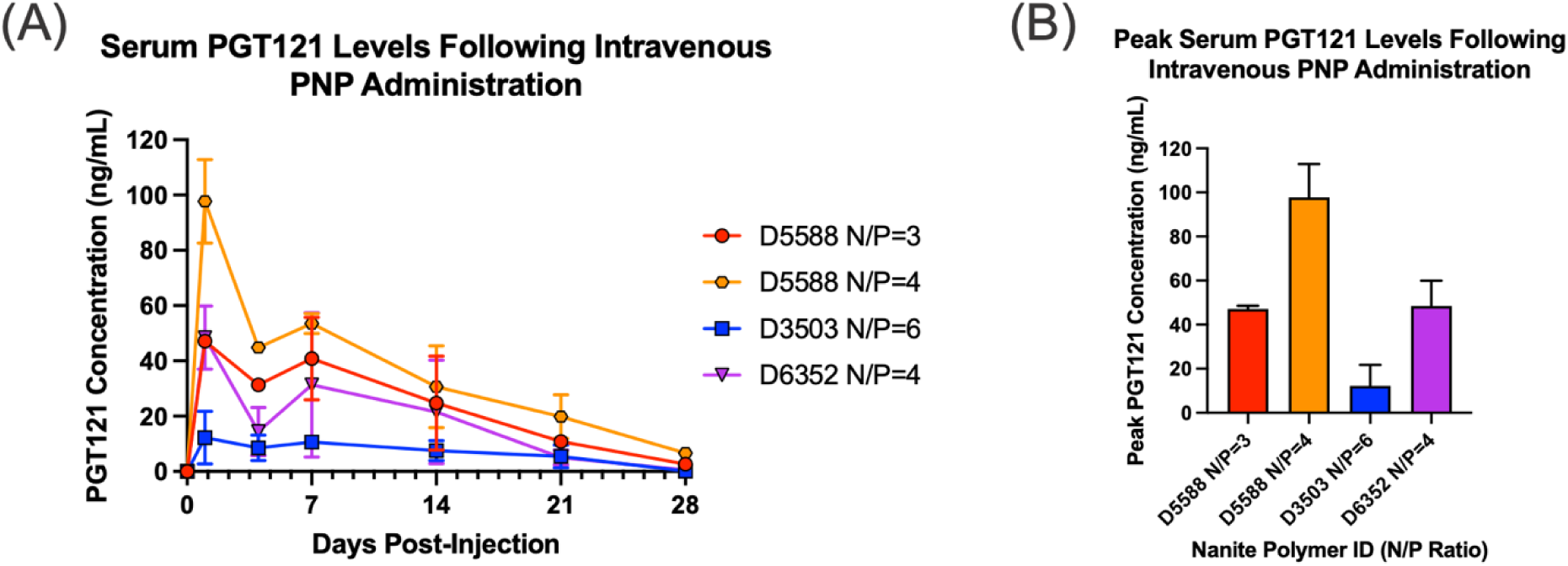
Serum antibody levels following intravenous delivery of Nanite PNPs. (A) Mouse serum PCT121 protein expression after i.v. administration of PNPs with PGT121 nanoplasmid at 1.8 mg/kg. (B) Peak serum PGT121 expression levels following intravenous administration from the experiments in Fig. 7A. n=2, error bars=SEM.

We next evaluated the delivery performance of our PNPs via an i.m. route of administration. Although targeting and biodistribution is localized with an i.m. injection, skeletal muscle cells have unique nanoparticle delivery challenges. We again took advantage of the RAG2-/-mouse model to carry out our i.m. injection experiments. PNPs were formulated and injected into the right tibialis anterior and the left gastrocnemius muscles for all animals. Serial blood draws were taken out to day 56 post-injection. Interestingly, we observed higher circulating PGT121 titers and different pharmacokinetic profiles and when compared to intravenous injection (Fig. 8A). When PNPs were delivered to mouse skeletal muscle, we achieved peak serum PGT121 levels between 0.400-1.100 μg/mL with an average peak expression time of 28-42 days post-injection (Fig. 8B). The antibody expression profiles were unique to the polymer, N/P ratio, and dose of nanoplasmid administered (Fig. 8C-F). The PGT121 expression levels were high (> 1.0 µg/mL), durable (sustained across many weeks), but also transient (decreased from peak levels). We observed no local toxicity at the site of injection nor any evidence of systemic toxicity (no decrease in body weights; data not shown). The ability of PNPs to tune antibody expression levels and profiles in vivo from DNA-encoded cargo based on polymer structure provides a foundation for further advances.

**Figure 8.**
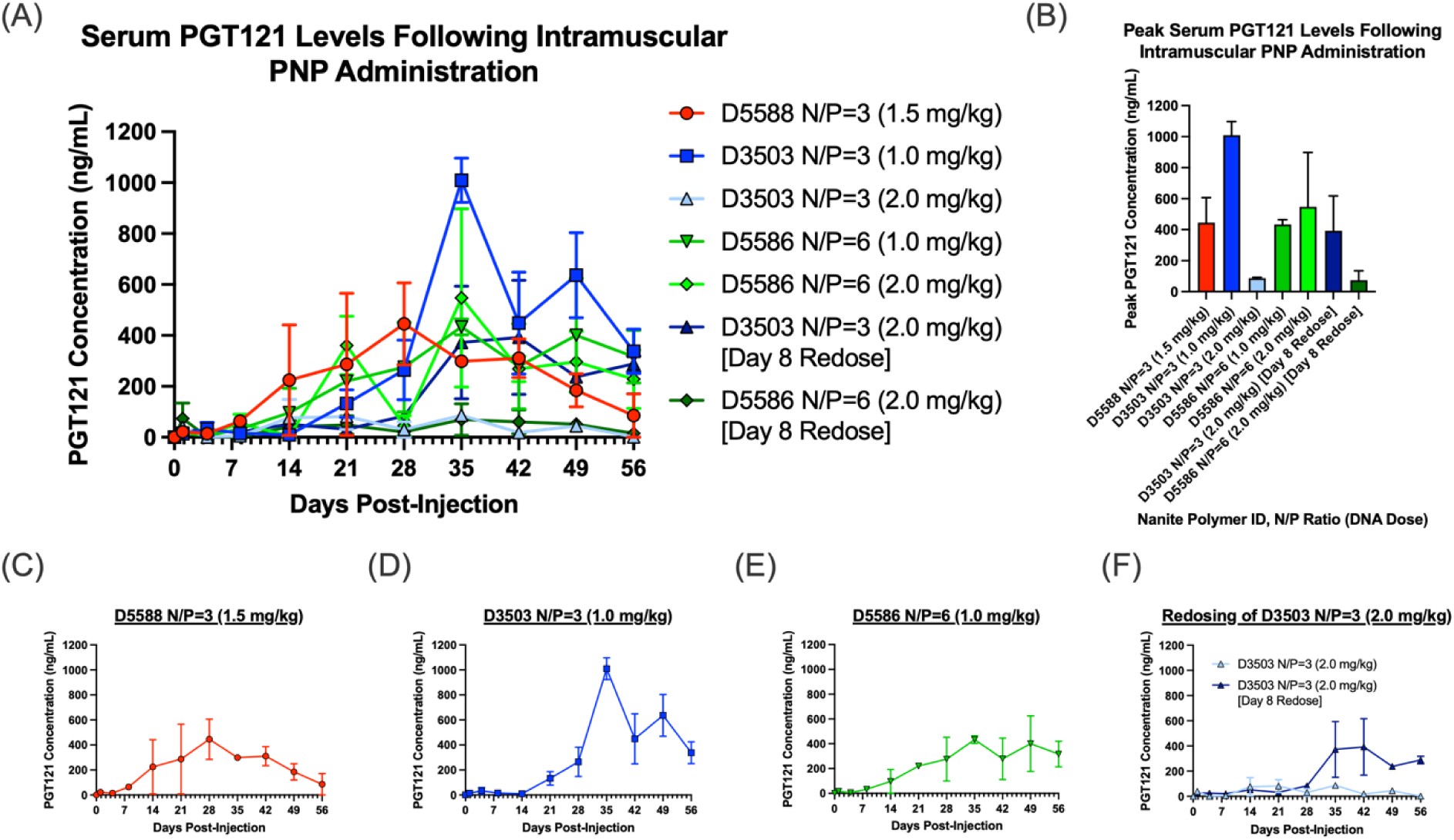
Serum antibody levels following i.m. delivery of PNPs. (A) ELISA readout of serum PGT121 antibody levels after i.m. PNP delivery of PGT121 nanoplasmid in RAG2-/-mice on Day 0. Nanite polymer, N/P ratio, and dose of nanoplasmid are denoted on the right for each condition. (B) Peak serum PGT121 expression levels following i.m. administration from the experiments in (A). (C) Focus on the time course of PGT121 serum levels from Nanite PNP D5588 shown in (A). (D) Focus on the time course of PGT121 serum levels from Nanite PNP D3503 shown in (A). (E) Focus on the time course of PGT121 serum levels from Nanite PNP D5586 shown in (A). (F) A single redosing of PNP D3503 at Day 8 (dark blue line) shows higher and more durable PGT121 serum levels when compared to a single dose of PNP 3503 (light blue line). n=2, Error bars=SEM.

Finally, we investigated the redosability of our PNPs, a challenge for many gene delivery vehicles, such as AAVs. Following an initial i.m. 2.0 mg/kg nanoplasmid PNP injection on day 0, D3503 PNPs were intramuscularly redosed with 2.0 mg/kg nanoplasmid at day 8 (Fig. 8F). We observed higher and more persistent PGT121 serum levels in the redosing group (dark blue line in Fig. 8F) when compared to a single dose on day 0 (light blue line in Fig. 8F). The ability to safely redose PNPs and improve circulating antibody titers in vivo is critical for genetically encoded antibody therapies where constant and predictable antibody levels are required. These data establish PNPs as safe, degradable, effective, and redosable gene delivery vehicles for DNA-encoded antibodies. Further, this DNA-encoded antibody PNP strategy could be applied to other clinically relevant secreted proteins.

## Discussion

In this work, we demonstrate the ability to use machine learning in combination with polymer chemistry to design PNPs capable of delivering a DNA-encoded anti-HIV antibody in vivo. These results suggest PNPs as a safe and effective DNA-delivery strategy that can achieve higher protein antibody levels at lower DNA doses than other delivery vehicles.^19^ PNP-delivery of a broadly neutralizing anti-HIV antibody could provide an alternative to daily antiretroviral therapy. It is particularly important to provide safe, inexpensive, easily manufacturable, and long-lasting treatment options to vulnerable patient populations in resource-limited geographies. We first characterized our machine learning-designed degradable PNPs in vitro to assess efficacy, toxicity, size, and stability (Figs. 5-6). The synthesized Nanite polymers formed small and stable PNPs with high transfection efficiency and low toxicity across a wide range of N/P ratios were therefore moved forward to in vivo testing. Next, we assayed the intravenous delivery capabilities of these PNPs. We observed strong serum PGT121 protein levels at 1-day post-transfection and a gradual decrease over time (Fig. 7A). These data show higher levels of protein expression when compared to other state-of-the-art DNA-delivery vehicles. As a more durable signal is needed for anti-HIV antibody therapies, we explored an alternative route of administration by direct injection of PNPs into the skeletal muscles of mice. Remarkably, i.m. delivery of Nanite PNPs provided safe, strong, durable, and transient protein levels with many conditions extending expression past the duration of the study. Importantly, these data show that polymer structure and therefore PNP structure, can be tuned to increase the serum antibody levels and duration of protein expression. We observed peak serum antibody levels one to two months after injection across nearly all conditions. PNPs containing polymer D3503 showed greater than 1.0 µg/mL peak protein expression levels with meaningful expression levels even at day 56 post-injection (Fig. 8E). Further, we showed that by delivering a DNA-encoded antibody using PNPs we can increase antibody levels and durability by redosing in vivo (Fig. 8F). PNPs containing polymer D3503 that were redosed at day 8 outperformed the single dose of the same condition (Fig. 8F). We noticed a general trend of lower dose and lower N/P ratio providing higher antibody levels when PNPs were delivered intramuscularly. These parameters, separate from polymer structure, provide different mechanisms by which we can optimize PNP in vivo delivery performance using machine learning techniques. Taken together, i.m. delivery of PGT121 nanoplasmid via Nanite polymers achieves meaningful levels of antibody expression over an extended duration, in a tunable manner. Additionally, these polymers are safe, degradable, and form small and stable particles. Further studies could be conducted to optimize these PNPs to provide even higher anti-HIV antibody expression levels and a more durable time course for direct use in a clinical setting. We view this work as part of a multi-pronged approach to lessen the HIV/AIDS disease burden on the most at risk patient populations. Finally, we emphasize that this strategy of delivering a DNA-encoded secreted protein via a safe and effective PNP in vivo could be applicable to a wide range of other disease modalities.

## Acknowledgement

We gratefully acknowledge funding from the Gates Foundation via Grant INV-066836.

